# The genetic architecture of hair colour in the UK population

**DOI:** 10.1101/320267

**Authors:** Michael D. Morgan, Erola Pairo-Castineira, Konrad Rawlik, Oriol Canela-Xandri, Jonathan Rees, David Sims, Albert Tenesa, Ian J. Jackson

**Author notes:** present address: Wellcome Trust Sanger Institute, Wellcome Genome Campus, Hinxton, Cambridge, CB10 1SA, UK. Joint first authors (Michael D. Morgan, Erola Pairo-Castineira).

## Abstract

We have extensively mapped the genes responsible for hair colour in the UK population. *MC1R* mutations are well established as the principal genetic cause of red hair colour, but with variable penetrance. We find variation at genes encoding its agonist (*POMC*), inverse agonist (*ASIP*) and other loci contribute to red hair and demonstrate epistasis between *MC1R* and some of these loci. Blonde hair is associated with over 200 loci, and we find a genetic continuum from black through dark and light brown to blonde. Many of the associated genes are involved in hair growth or texture, emphasising the cellular connections between keratinocytes and melanocytes in the determination of hair colour.

## Introduction

Natural hair colour within European populations is strikingly variable, and is a complex genetic trait that is impacted little by non-genetic, environmental factors. Furthermore, hair colour is largely determined by only a few well-characterised cell types: the melanocytes where the melanin pigment is made, the keratinocytes of the hair to which the pigment is transferred, and fibroblasts of the dermal papilla, which signal to and regulate the melanocytes. It is thus an excellent model system to explore genetic and cellular interactions in development and homeostasis. Hair colour variation is partially correlated with skin and eye colour variation, reflecting differences in cellular interaction in different tissues (1, 2).

Several studies have examined the genetic basis of hair colour variation. Red hair is well established as being associated with coding variation in the *MC1R* gene (3) (4). Less well known is the observation that most of these variants are only partially penetrant, and some of them have very low penetrance indeed (5). Other genetic factors must be interacting with *MC1R* to modify the penetrance of these variants. MC1R is a G protein coupled receptor, expressed on the surface of skin and hair melanocytes. Binding of the MC1R cognate ligand, α-melanocyte stimulating hormone (α-MSH), induces a melanogenic cascade resulting in the production of dark eumelanin. This is packaged into vesicles, termed melanosomes, for transport to epidermal keratinocytes where it provides protection against ultraviolet radiation. The cellular trafficking of melanosomes to keratinocytes in the hair follicle additionally gives colour to the growing hair. Loss of MC1R signalling in many vertebrate species results in the inability of the melanocytes to produce eumelanin that instead default to synthesising phaeomelanin, a red or yellow pigment. MC1R has a second ligand, an inverse agonist, agouti signalling protein (ASIP) (6). Overexpression of ASP, in mice for example, leads to synthesis of only yellow phaeomelanin, even in the presence of a functional MC1R and α-MSH (7). Previous studies have noted an association between the *ASIP* locus and red hair in humans, but the nature of the association has not been explored (8).

Until recently, genome-wide association studies (GWAS) identified only a small number of loci are associated with blonde hair, compared to black and brown (9), (10), (11) (12). Each of these studies identified between 4 and 8 genes, with a total of 11 genes associated with hair colour differences. Most of these loci have been previously described as causing coat colour variation in mice (*MC1R*, *ASIP*, *OCA2*, *SLC45A2*, *KITLG*, *TYR*, *TYRP1*, *EDNRB*), zebrafish (*SLC24A5*) and humans (*TPCN2*, *IRF4*). However, during preparation of this manuscript a GWAS was reported identifying over 100 loci contributing to hair colour (13). This report however, did not distinguish between genes affecting different hair colours.

Hysi et al (13) analysed a subset of participants in a very large population health cohort of British individuals, UKBiobank in addition to a similar number of individuals from 23 and Me, totalling 290,891. We report here the analysis of the majority of UK Biobank, a total of almost 350,000 subjects. By performing genome-wide analyses across hair colours, we have discovered novel variation in and around *MC1R* that contributes to red hair. Furthermore we identify 7 additional loci that contribute to red hair, including at *ASIP*, where an eQTL shows epistatic interactions with the poorly penetrant *MC1R* variants. Additional epistatic interactions are seen between *MC1R* and the *HERC2/OCA2* locus and with *PKHD1*. In addition we identify more than 200 loci associated with multiple hair colours on the spectrum of blond to black. Notably, we find that many of the associated genes are not involved in melanocyte biology *per se*, but are rather involved in hair growth or texture. This highlights the importance of the melanocyte-keratinocyte interactions in the determination of hair pigmentation and the impact of hair shape on colour perception.

## Results

Participants in UK Biobank responded to the question “what is your natural hair colour” with one of 6 possible answers. We used only self-reported, white British individuals, confirmed by genotype (14). In addition, of the individuals who were 3^rd^ degree relatives (first cousins) or closer, identified by genotyping (14), only one of any related group was analysed. This left 343,234 participants with hair colours: red n=15731 (4.6%), blonde n=39397 (11.5%), light brown n=141414 (41.2%), dark brown n=127980 (37.3%), black n=14526 (4.2%), other n=4186 (1.2%).

Genotypes were determined by UK Biobank for more than 800,000 SNPs and indels, which were directly assayed, and an additional ~40M imputed using the Haplotype Reference Consortium panel (14).

### Red hair colour and MC1R

As *MC1R* variation is well established as contributing to the genetic basis for red hair, we first analysed coding variation in this gene. We find that 86.3% of red-haired individuals carry two non-synonymous, frameshift or nonsense variants (homozygous or compound heterozygous) and 11.7% carry a single variant. ~1% have three variants and less than 1% have no MC1R coding variants detected. The cases of red hair with only one or no variants (similar to that seen in a study of an Australian cohort (5) may be explained by a) rare coding variant alleles not genotyped or imputed in this study, b) extragenic variation affecting *MC1R* expression c) dominant action of specific alleles d) variation in other genes in the same or a parallel pathway or e) misreporting of hair colour. In our genotyped and imputed panel of variants, we identified two additional coding variants, rs368507952 (R306H) and rs200000734 (R213W) and a variant, rs3212379, located 120bp 5’ of the transcription start site of *MC1R*, which is a candidate transcriptional regulatory variant. Including these in our analysis increases the proportion of red-haired individuals with two *MC1R* alleles to 92%, whilst only 6.3% carry a single allele.

It is well established that different *MC1R* coding variants have different penetrance with respect to red hair (termed “R” and “r” for high and low penetrance) (5). With this very large cohort we are able to more precisely quantify the degree of penetrance of each allele, whether as homozygotes or in combination with any other allele. (Figure 1, Supplementary Table 1). Similar to others we find that penetrance of missense variants ranges from less than 1% as homozygotes (V60L, V92M) to over 90% (D294H).

**Figure 1.**
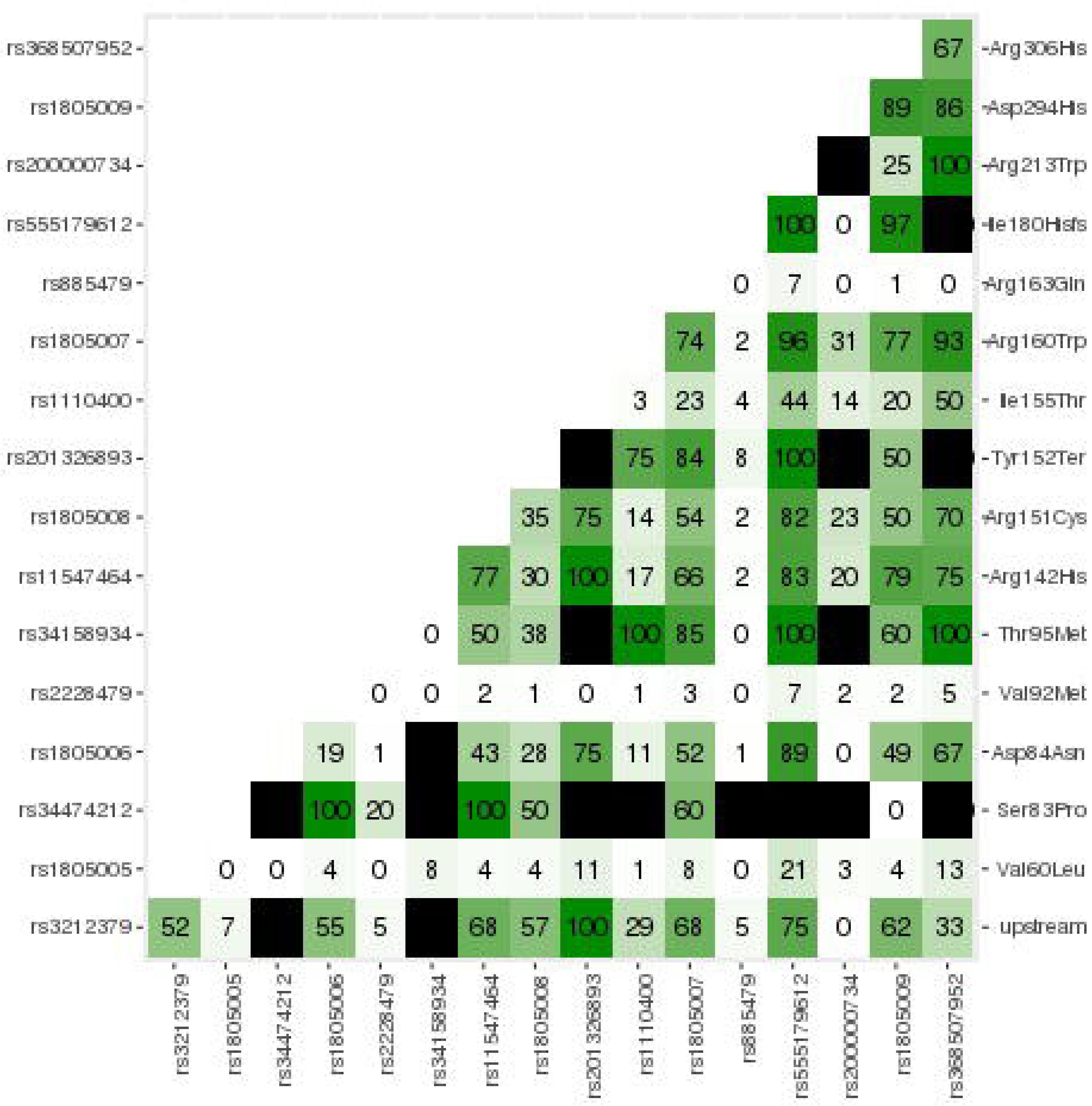
Penetrance matrix of *MC1R* coding variants. Combinations of all coding variants plus the non-coding variant rs3212379, located close to the 5’ end of *MC1R*. Cells indicate the % of the genotype with red hair, rounded to whole numbers. Cells filled in black have no data. Full data is in Supplementary Table 1.

**Table 1:** Enrichment for cell type specific annotations. GWAS loci from all hair colours. P-values are adjusted using the Bnejamini_Hochberg method. The Δenrichment column represent the change in enrichment (nsnpOverlap/allsSnps) between our dataset and the median of the 10000 permutations.

| Cell Type | Annotation | BH adjusted p-value | Δenrichment |
| --- | --- | --- | --- |
| Melanocyte | H3K27ac | 0.4930 | 0.00317 |
|  | H3K27me3 | 0.0726 | 0.00460 |
|  | H3K36me3 | 1 | -0.00576 |
|  | H3K4me1 | 1 | -0.02505 |
|  | H3K4me3 | 0.00054 | 0.01526 |
|  | H3K9me3 | 0.00054 | 0.00921 |
|  | DHS | 1 | -0.01382 |
|  | TSS | 1 | -0.00259 |
|  | MITF | 0.0009 | 0.01497 |
| Keratinocyte | H3K27ac | 1 | 0 |
|  | H3K27me3 | 0.0453 | 0.00749 |
|  | H3K36me3 | 1 | -0.02160 |
|  | H3K4me1 | 0.27196 | 0.00605 |
|  | H3K4me3 | 0 | 0.02418 |
|  | H3K9me3 | 1.0 | -0.01785 |
|  | TSS | 1 | -0.00432 |
| Fibroblast | H3K27ac | 0.484875 | 0.00346 |
|  | H3K27me3 | 0 | 0.02246 |
|  | H3K36me3 | 0.26352 | 0.00461 |
|  | H3K4me1 | 1 | -0.01180 |
|  | H3K9me3 | 0.0038 | 0.00720 |
|  | DHS | 1 | -0.02332 |
| Iris pigment | TSS | 0 | 0.00519 |
| Dermal fibroblast | TSS | 1 | -0.00259 |
| Skin Fibroblast | TSS | 1 | -0.00086 |
| Skin Tissue | TSS | 1 | -0.00375 |
| Embryoid melanocytic induction | TSS | 1 | -0.00230 |
| Hermes3A | MITF | 1 | -0.00374 |

To identify the genetic variation that might account for the unexplained cases of red hair, and that might influence the penetrance of the *MC1R* variants, we performed a GWAS on the red haired vs. black and brown hair individuals combined. Accounting for genetic structure within the UKBiobank by inclusion of the first 15 genetic principal components adequately controlled the genomic inflation in our analysis (λ_GC_ 1.018). The strongest association with red hair is located around the *MC1R* gene on human chromosome 16 (Figure 2a Supplementary Table 2, Supplementary Figure 1), which fits with the expectation that this locus is the principal genetic factor determining red hair colour. We find that the strongest signal of association in the region of *MC1R* (rs34357723; OR 9.59, p<2.25×10^−308^) does not originate from any observed amino acid changes, nor are there any known non-synonymous *MC1R* mutations in moderate to strong linkage disequilibrium (1000 Genomes EUR LD r^2^≥0.5). Indeed this association is with a SNP located some 97kb from the 5’ end of *MC1R*, and remains significant even after adjusting for all coding variants in MC1R. As we know that multiple *MC1R* alleles affect red hair colour, we performed step-wise conditional association testing, and identified 31 additional association signals in this region at genome-wide levels of statistical significance (p≤5×10^−8^), altering the odds of having red hair compared to brown and black hair (Supplementary Table 2). Only 10 association signals can be directly attributed to amino acid changes, nonsense or frameshift mutations within the *MC1R* coding region. Included in these are the two missense variants rs368507952 (R306H) and rs200000734 (R213W) noted above, not previously associated with red hair colour.

**Figure 2.**
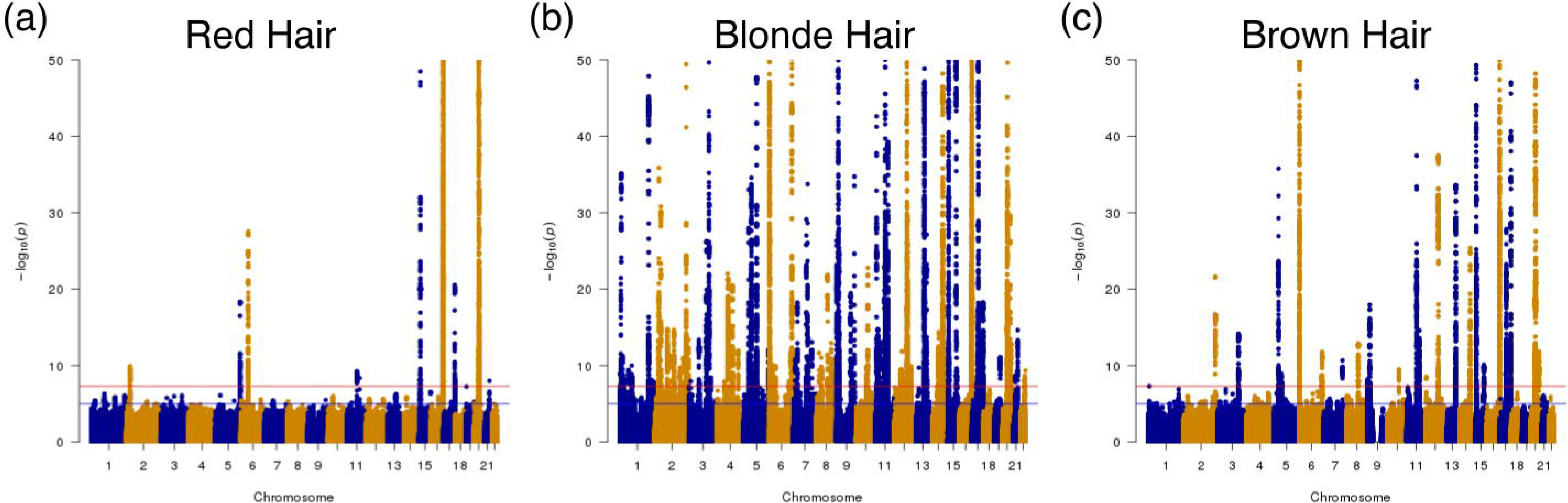
Manhattan plots of GWAS data. Data plotted for (a) red hair versus black plus brown hair, (b) blonde hair versus black plus brown hair, (c) brown hair versus black hair. Points are truncated at –log10(p) = 50 for clarity

**Table 2.** Phenotype of mouse mutants at candidate genes. Skin/Skin Appendages refers to all skin phenotypes (skin, hair, teeth, sweat glands, mammary glands) except pigmentation. Genes in bold are orthologues of genes identified that affect hair shape variation

| Skin/Skin Appendage Phenotype |  |  | Pigmentation Phenotype |  |
| --- | --- | --- | --- | --- |
| Alx4 | Frem2 | Msx2 | Dct | Pax3 |
| Areg | Grm5 | Ndufs4 | Edn3 | Plk2 |
| Asb | Hdac4 | Ovol1 | Ednrb | Plxnb2 |
| Bmp7 | **Hoxc13** | **Padi3** | Frem2 | Slc45a2 |
| Ccnd1 | Il6 | Ppm1a | Kitl | Tyr |
| Cdkal1 | Kat6a | Rspo2 | Mc1r | Tyrp1 |
| En1 | Krt33a | Sp6 | Mitf | Xpa |
| **Errfi1** | **Krtap17-1** | Syne2 | Mkln1 | Zmiz1 |
| Fgf5 | Lef1 | Twist2 | Oca2 |  |
| Fosl1 | **Lgr4** | Xpa |  |  |
| Foxe1 | Lhx2 |  |  |  |
| **Fras1** | Map3k1 |  |  |  |

In addition to these 10 coding variants, we find 21 associations beyond the *MC1R* coding region at distances from 120bp 5’ (e.g. rs3212379), to up a megabase both 3’ and 5’. These distal associations cannot be attributed to LD with *MC1R* variation and have been observed in other studies ((1), (10)). Potentially these variants affect long range regulatory elements of *MC1R*.

### Additional red hair colour associated loci

In addition to the associations around *MC1R* on chromosome 16, we observe 7 additional associations at genome-wide significance (Suppl Table 2). Statistical fine mapping of causal SNPs (PICS) (15) in some cases indicated a single likely casual variant, whilst in others one of more than 50 variants could be the causal SNP. We find a previously unreported association at r276645354. This variant lies less than 2kb from the transcriptional start of *POMC*, which encodes α-MSH, the agonist of MC1R. A single variant in an intron of *RALY*, located 5’ of *ASIP*, the gene encoding the inverse agonist of MC1R, was associated with red hair. This variant, rs6059655, is also an expression QTL (eQTL) for *ASIP* expression in skin, with the red-hair associated allele showing higher mean expression levels (16) (Gtex website) (Figure 3). We suggest that variants which increase *ASIP* expression in the skin or hair follicles lead to greater competition with α-MSH for melanocyte MC1R binding, dampening melanogenic induction, and increasing the pheomelanin in melanocytes.

**Figure 3.**
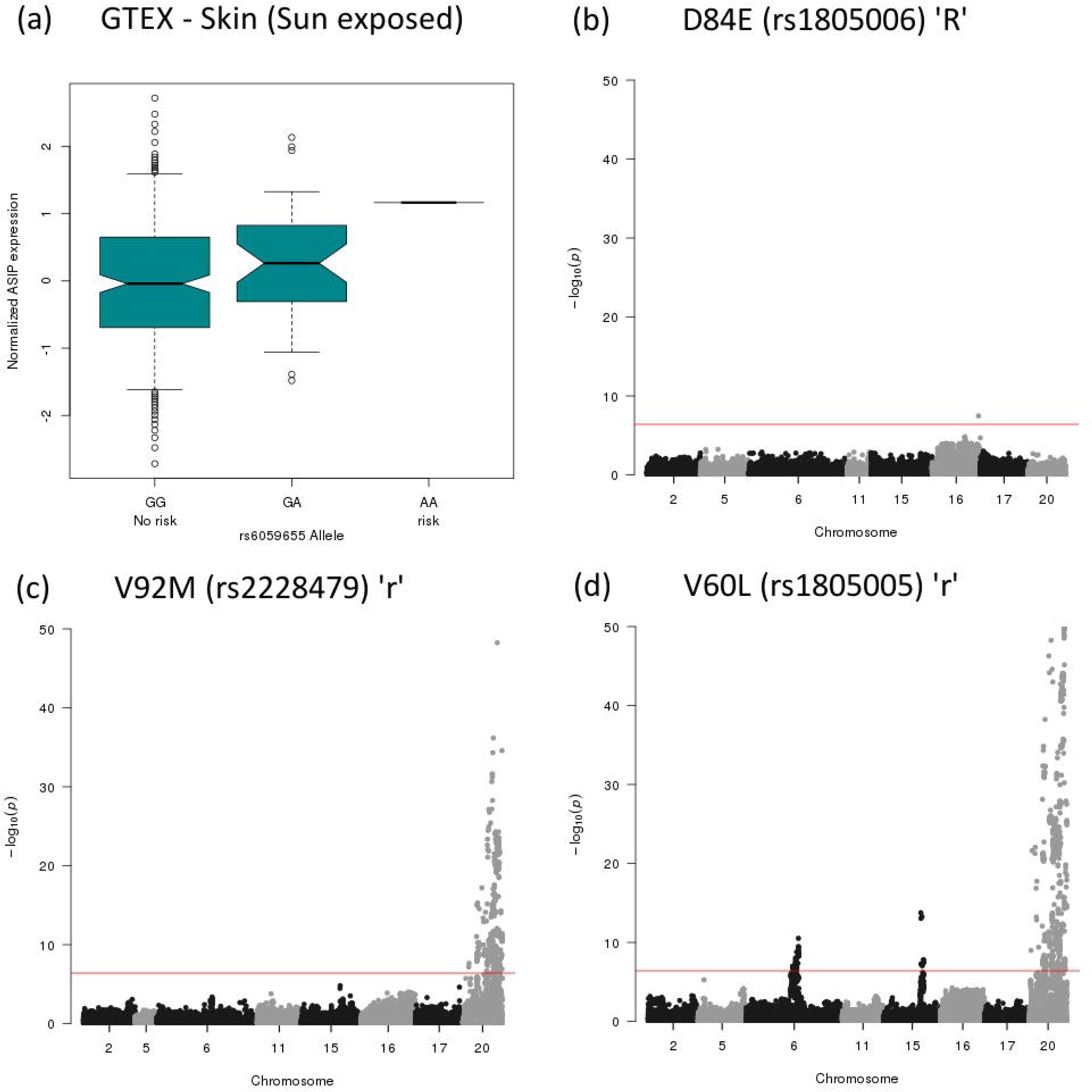
Gene expression variation at ASIP and epistatic interactions with MC1R variants. (a) Gene expression data from GTex of *ASIP* in sun-exposed skin, ordered by genotype at rs6059655, and normalised to the homozygous no-risk genotype (GG). (b-d) Epistatic interactions between *MC1R* coding variants and other red-hair associated loci. (b) The high penetrant allele D84E shows no *trans* interactions (c) the low penetrant allele V92M shows interactions at *ASIP* (d) the low penetrant allele V60L shows interactions with *ASIP*, *HERC2/OCA2* and *PKHD1*

We find a variant in *HERC2* associated with a decreased probability of red hair. It is well established that variants in *HERC2* alter transcription of the neighbouring pigmentation gene *OCA2*, and are additionally associated with blue eyes and blonde hair colour (10, 17, 18), (19, 20).

Association is also seen in the *TSPAN10* gene, also known as oculospanin, which is highly expressed in melanocytes and retinal pigment epithelium. The lead SNP lies in strong LD (r^2^=0.995) with a non-synonymous variant (rs6420484; Y177C) affecting a conserved amino acid. This association signal is also in moderate LD (r^2^ ~ 0.4, D’ ≥ 0.95) with a previously reported association with increased hue-saturation of eye colour, which corresponds to darker eyes, in a Dutch cohort (21). Previous targeted knockdown of the murine *Tspan10* mRNA resulted in reduced melanocyte migration in a trans-well migration assay (22), indicating this gene may be a good functional candidate as a novel hair colour gene.

### Epistasis between MC1R alleles other loci

Detecting epistasis in complex traits is challenging. Epistatic effects are believed to be much smaller than main effects, which are typically already very small in the case of polygenic traits. However, due to the large effects of some genetic variants on hair colour this might be a more tractable model to detect epistasis. We tested all associated genetic variants from our analysis of red hair colour against each of the *MC1R* coding variants, including the known ‘R’ and ‘r’ *MC1R* red hair colour alleles, by constructing a logistic regression model whilst correcting for relevant covariates (see Methods). At a p-value of 3.9×10^−7^ (i.e. 0.05/128205, the number of variants tested) we found consistent epistasis signals between *MC1R* variation and a ±1.5MB region surrounding rs6059655, which is the hair colour associated *ASIP* eQTL SNP (Figure 3, Supplementary Figure 2). We detect epistasis between rs1805005 (V60L) and rs1805008 (R160W) and the *HERC2/OCA2* region. It has been noted previously that *OCA2* variation affects the penetrance of the weaker red hair alleles of *MC1R* (5). Finally, we also can detect epistasis between V60L and *PKHD1* on chromosome 6. The magnitude of the UK Biobank cohort has allowed the identification of hitherto unknown epistatic interactions.

### Genome-wide association analysis of blonde hair colour

Whilst red hair is essentially a Mendelian trait modified by additional loci, the genetic architecture of blonde hair colour is concordant with a polygenic trait. We performed a genome-wide association analysis comparing blonde to combined brown and black hair colour. Following conditional association testing to uncover additional signals of association at a number of loci, we discover 213 loci associated with blonde hair colour (Figure 1b, Supplementary Table 3, Supplementary Figure 1b). In many cases multiple signals of association are found close to the same genes. This could be a result of multiple, independent associations (as is the case for *MC1R*, for example). Alternatively some or all may each be correlated with the same variant that has been neither genotyped nor imputed.. Many signals of association are close to, or within, previously known pigmentation genes from both human and model organism studies. These allelic effects span a spectrum of odds ratios and minor allele frequencies consistent with many other phenotypes with an underlying polygenic architecture (23, 24) (Figure 4a). We are able to statistically refine ~1/3 of these association signals to a single candidate variant which include multiple non-synonymous variants. Several variants notably stand out, which have been previously associated with a variety of pigmentation related traits in humans (including *SLC24A4, HERC2/OCA2, SLC45A2, TYR, TYRP1, EDNRB*), some of which have been linked to alterations in transcriptional regulation (*IRF4* & *KITLG*) (25, 26).

**Figure 4.**
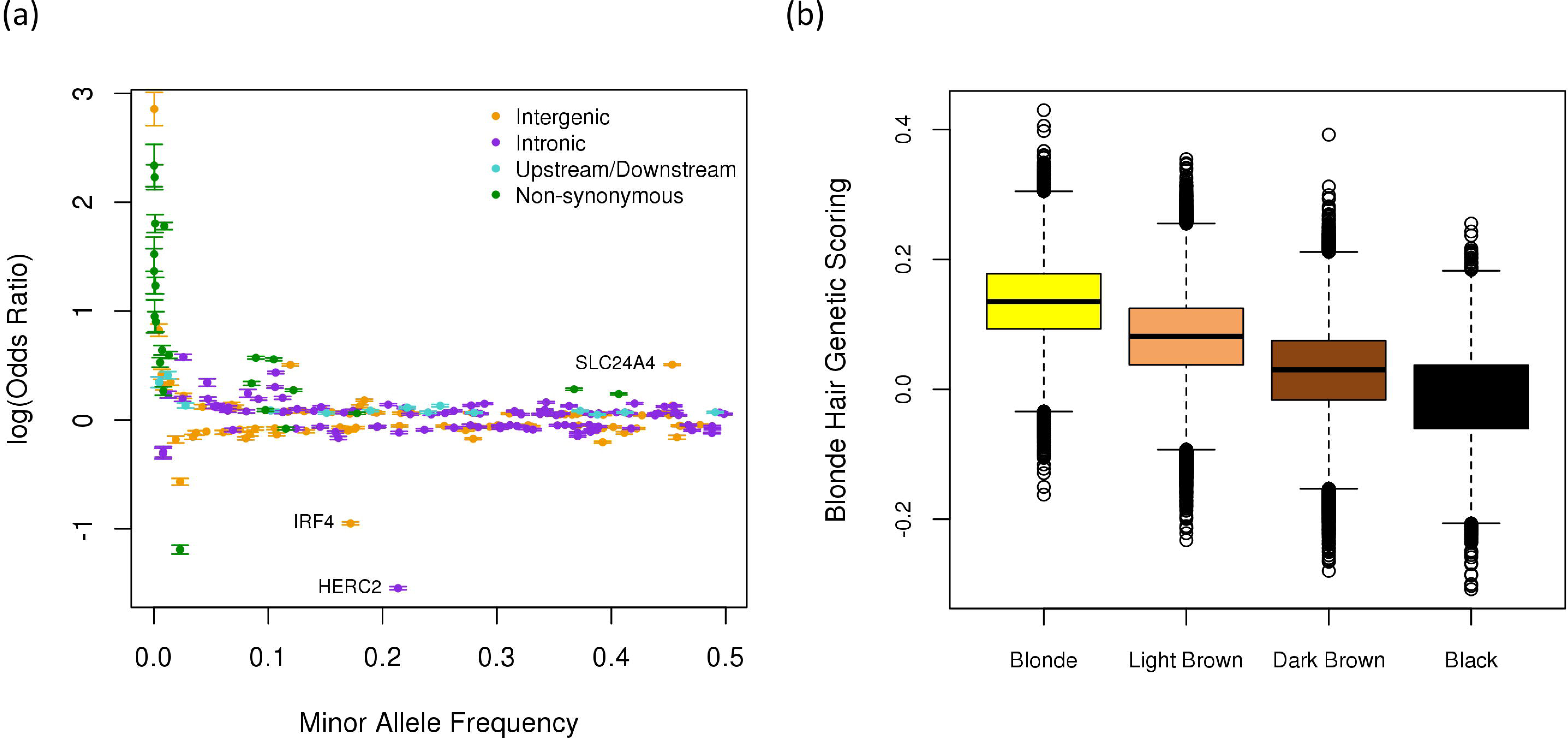
Odds Ratio and Minor Allele frequency for Blonde Hair and Polygenic Phenotype Scores. (a) Plot of minor allele frequency of blonde hair associated variants versus log of the odds ratio for blonde hair. Variants are colour-coded for annotation; intergenic (yellow), intronic (purple), 2kb upstream or 500bp downstream (cyan), non-synonymous coding (green). (b) Genetic scores derived from all lead variants from blonde versus brown plus black hair colour, assuming an additive genetic model.

### Genome-wide association analysis of brown hair colour

We hypothesised that hair colour may lie on a continuous genetic spectrum. Thus, we might expect to observe a subset of the blonde associated variants associated with brown hair. Following both primary and conditional analyses we find 56 loci associated with brown versus black hair (Supplementary Table 4; Figure 1c, Supplementary Figure 1c), 28 of which are the same associated variant with blond hair, and with the same direction of effect. Of the remaining loci 23 identify the same genes seen in our blonde hair analysis. One of the novel genes, *PIGU*, is not seen explicitly associated with blonde hair, although we observe an association with another member of the same gene family, *PIGV*, suggesting that paralogous genes may be associated with hair colour differences.

### Polygenic phenotype scoring

To test the hypothesis that the genetic basis of hair colour is polygenic and that hair colour falls on a continuum as a genetic trait, we constructed a polygenic score for hair colour. We constructed a blonde hair colour polygenic phenotype score by taking the variants that reached genome-wide significance in the blonde vs. brown and black hair colour analysis (5×10^−8^), as a linear combination of the allele-weighted regularized logistic regression coefficients. We found that self-reported black, dark brown, light brown and blond hair lie on an approximately linear spectrum (Figure 4b). We confirmed the same pattern across hair colours in two groups of individuals excluded from all previous analyses; related individuals (Supplementary Figure 3a), and individuals with European, but non-British ancestry (Supplementary Figure 3b).

### eQTL

In order to aid the interpretation of our GWAS, and identify functional hypotheses, we tested loci for statistical colocalisation with eQTL signals from skin-biopsies in the GTex (16), (Supplementary Tables 5-10) and TwinsUK cohorts (27), (Supplementary Tables 11-13). We were able to link 37 loci with *cis* eQTLs with high probability (posterior probability >0.8). Among the variants with the highest probability are at *RALY*, upstream of *ASIP* as noted above and in the first intron of *TSPAN10* for red hair. Most of the eQTLs are associated with gene expression at considerable distance, and often with several genes. Curiously also among the most significant eQTLs, across all three datasets, are several missense variants in *MC1R*, which are independently linked to expression changes in multiple genes located up to several hundred kb from *MC1R*, reflecting again the unusual behaviour of this segment of the genome. Whist the colocalisation of hair colour association signals with skin tissue *cis*-eQTLs may appear promising they are at best a strong indication of biological effect, and will require extensive further hypothesis testing to establish any role in determining pigmentation.

### Hair colour loci are enriched for regulatory features in pigmentation cell types

To understand the transcriptional regulatory mechanisms that might underpin the observed genetics associations with hair colour we examined the potential for these variants to affect the chromatin landscape in cell types relevant to pigmentation. Specifically, we tested histone tail modifications associated with gene activation or repression and with chromatin accessibility (DNase I hypersensitive sites) in melanocytes, keratinocytes, fibroblasts and other cells (Table 1). Additionally, the proximity of several association signals to core promoter regions raises the possibility of alterations to transcriptional start sites (TSS) and pigmentation cell-specific regulatory factors, i.e. the melanogenesis master regulator MITF. Using a permutation-based approach (GoShifter) (28), we tested each annotation in each cell type where data were available (Table 1). We find statistical evidence of enrichment of pigmentation associated genetic variation overlapping histone marks of both gene activation (H3K4me3) in melanocytes and repression (H3K9me3, H3K27me3) in melanocytes, fibroblasts and keratinocytes. In addition there is enrichment of MITF binding sites in melanocytes and TSS in iris pigmentation cells. These associations give strong support to the notion that we are able to identify functional elements altered by genetic variation.

### Enrichment for Skin and Hair Genes

To further aid the interpretation of our GWAS findings, and identify shared biological pathways related to pigmentation determination, we took all of the blonde hair lead SNPs overlapping genic regions extending 2kb up stream of the TSS, and 500bp downstream of the 3’ end. If no genic region overlapped the lead SNP, then we used the two closest genes within 500kb (Supplementary Table 14). These candidate genes were then used as input to test for enrichment in known pigmentation phenotypes, utilising the MouseMine database (29). We identified ~200 orthologous mouse genes in the database, which we analysed for site of expression and mutational phenotypes. Of the 172 genes with expression data, 89 were expressed in the skin (P = 1.3 × 10^−9^) (Supplementary Table 15). 132 genes had mouse mutant phenotype data, and of these, 50 had an integument phenotype (affecting the skin and skin appendages) (P=5.2×10^−7^). Not surprisingly, 18 of these affected pigmentation, but we unexpectedly find that 70% affect primarily skin, hair or other skin appendages rather than pigmentation (Table 2)

Follicular melanocytes, keratinocytes and dermal papilla cells have mutual interactions; the dermal papilla signals to melanocytes with ASIP, the melanocytes transfer melanin granules into the keratinocytes. Perturbations of these interactions could affect the amount and type of melanin delivered to the hair. Furthermore, variation in growth rate could impact the effectiveness of melanin transfer. Recent GWAS have identified 14 loci associated with hair shape variation (30). Remarkably, we have identified 7 of these, *ERRFI1*, *FRAS1*, *HOXC13*, *PADI3*, *KRTAP*, *PEX14* and *LGR4* as affecting blonde/non-blonde hair colour (P=1×10^−11^, Fishers exact test). In addition, the refractive and reflective properties of individual hairs may affect perceived colour (31) and there is evidence that different coloured hairs have different morphology. Vaughn et al have demonstrated a strong inverse correlation between the lightness of hair colour and the diameter of the shaft; blonde hair is thinner than dark (32). In summary the very large dataset provided by UK Biobank has enabled us to dissect the complex genetic nature of hair colour. This forms the foundation for functional analysis linking genetic variation to phenotype, and exploring the cellular interactions between melanocytes and other cells in the hair follicle.

## Methods

Study individuals were derived from the UK Biobank cohort that consists of 502,655 individuals aged between 40 and 69 years at recruitment, ascertained from 22 centres across the UK between 2006 and 2010. From these, we analysed 343,234 unrelated individuals, which reported their background as “British” and with similar ancestral backgrounds based on PCA (14).

Imputed and genotyped autosomal variants were included in analysis, with a Hardy-Weinberg equilibrium test p-value > 10^−10^, a call rate > 0.95 in unrelated white British individuals, and with a UKBB score > 0.8.

Self-reported hair colour (before greying occured) for UK Biobank participants was selected from one of eight possible categories: “Blonde”, “Red”, “Light brown”, “Dark brown”, “Black”, “Other”, “Prefer not to answer”. The number of individuals for each hair colour ranged from 141,414 for light brown to 14,526 for black.

To test for the association between hair colour and genotype we fitted a logistic regression model adjusting for population structure, using the first 15 axes of variation from the PCA (14), and genotyping batch.

Conditional analyses used the same multivariate logistic regression model, with the addition of the lead genetic variants as covariates. A forward selection variable strategy was followed. That is, we kept adding new variants to the model one by one (starting from the most significant one) until we added one that did not reach a significant level of 5×10^−8^. PLINK v1.9. was used for the regression analysis and Manhattan and Q-Q plots were generated using the R package *qqman* (33) and *ggplot2*.

We implemented the probabilistic inference of causal SNPs (PICS) (34) which takes into account the strength of association of the lead genetic variant and the LD structure in the fine-mapping interval to calculate the posterior probability that a particular genetic variant is the causal variant given that another variant is the lead variant (see Supplementary Methods for details).

Variants within 3 Mb of the lead variant (±1.5Mb either direction) for each signal of association were selected for epistasis analysis with *MC1R* variants for red hair colour. Epistasis was tested with a likelihood ratio test, comparing a full logistic model that includes an interaction term with a reduced logistic model without the interaction. Epistasis testing was performed using *cassi* (see URLs) and included the same covariates as our original genome-wide and conditional analyses (i.e. first 15 PCs and genotyping batch). Due to complex LD structure around the *MC1R* region it was necessary to remove variants in LD (r^2^ ≥ 0.1) with the target variant in each case. Additionally, we corrected for the most significant variants in the *MC1R* region, to correct for the influence of variants in very low LD.

In order to calculate the hair colour genetic score we performed least absolute shrinkage and selection operator (LASSO). A penalized logistic regression model was created for blonde colour, including all lead variants for blonde vs. black and brown hair colour. The parameter λ was selected using the most regularized model from a 10 fold cross validation, within 1 standard error of the minimum using the *glmnet* package in R. The regularized effect estimates were used to create genetic scores for blonde hair colour, assuming an additive genetic model.

Epigenetic marks (H3K4me1, H3K4me3, H3K27ac, H3K17me3, H3K36me3, H3K9me3 and DNase I hypersensitive sites) from the Roadmap Epigenome and ENCODE project were downloaded for primary melanocytes, keratinocytes, and fibroblasts. Transcription start sites were downloaded from FANTOM5 for primary melanocyte, keratinocyte and other pigmentation cell types. MITF ChIP-seq data derived from primary melanocytes and the melanocytic cell line Hermes3A were downloaded from NCBI Gene Expression Omnibus (GEO) and Short Read Archive (SRA). Short sequencing reads were extracted using *fastq-dump* and assessed for quality using *fastQC* (see URLs). Reads were processed in order to remove adaptor contamination using *trimmomatic* and then aligned to GRCh37/hg19 using BWA-aln (35). As there are no replicates in the ChIP-seq data, peaks were called with MACS2 (36) with an FDR 1%.

For each cell type and annotation where more than one sample existed, intervals were merged using *Bedtools* (37). To test for genomic enrichment we used a method that tests for the local enrichment of lead variants and those in LD with genomic features by randomly shifting annotations to generate a null distribution; implemented in *GoShifter* (38). LD values calculated from the 1000 Genomes phase 3 EUR reference panel were used in conjunction with the lead variant from each independent signal of association (after conditional analyses). *GoShifter* was run for 10,000 permutations, and enrichment p-values were adjusted for multiple testing using the Benjamini & Hochberg procedure implemented in the R function *p.adjust* with a 5% FDR.

To test for the potential of non-coding association signals to impact on gene regulation we utilised expression quantitative trait loci (eQTL) summary statistics from the GTex (39, 40) and MuTHER (41, 42) studies in tissues relevant to pigmentation (MuTHER; skin biopsies, GTex; sun-exposed and sun not-exposed skin tissue). Summary statistics for genes that fell within 3Mb intervals of the lead SNP of each association signal were extracted. We wished to test the explicit hypothesis that the same causal variants underlie each eQTL and our association signal for each hair colour (H_A_). Subsequently the null hypothesis is a compound null of no association for either trait (H_01_), and the combinations of overlapping but independent eQTL and trait associations (H_02_) and an association in one trait/eQTL but not the other (H_03_ & H_04_). A Bayesian testing framework that tests the set of hypotheses relating to those defined above (H_01-4_ & H_A_) was developed by Giambartolomei *et al*, and is implemented in the R package *coloc* (43). Subsequently we report the posterior probability for each association signal-gene eQTL signal and hypotheses H_02-04_ (PP1, PP2 & PP3) and the alternative hypothesis H_A_ (PP4).

URLs
*Gtex*: https://www.gtexportal.org/ *Twins UK*: http://www.muther.ac.uk/ *Cassi*: http://www.staff.ncl.ac.uk/richard.howey/cassi/ *SRAtoolkit:* https://github.com/ncbi/sra-tools *fastqQC*: https://www.bioinformatics.babraham.ac.uk/projects/fastqc/

## Acknowledgements

This work was carried out under UK Biobank study number 7206. It was funded by MRC core support to the Human Genetics Unit and to the Computational Genomics Analysis and Training programme through grant G1000902 and by BBSRC funding through Strategic Grant funding to the Roslin Institute. We would like to thank Sebastian Luna-Valero for extensive systems admin support, and the other members of the CGAT programme for numerous robust and constructive discussions.

## Supplementary Materials and Methods

### 1. Study participants

Study individuals were derived from the UK Biobank cohort that consists of 502,655 individuals aged between 40 and 69 years at recruitment, ascertained from 22 centres across the UK between 2006 and 2010. From these, we analysed 343,234 unrelated individuals, which reported their background as “British” and with similar ancestral backgrounds according to PCA^1^

### 2. Genotype Quality Control

Variants included in the analysis were autosomal SNPs present in the HRC imputation panel, with a p-value for the Hardy-Weinberg equilibrium > 10^−10^, a call rate > 0.95 in unrelated white British individuals, a UKBB score > 0.9 and a MAF > 10^−4^.

The number of SNPs analysed after quality control is 9,154,080

### 3. Phenotype quality Control

Self-reported hair colour data from UK Biobank cohort was used, consisting of eight categories: “Blonde”, “Red”, “Light brown”, “Dark brown”, “Black”, “Other”, “Prefer not to answer”, “Do not know”, The number of individuals in each category are: Red: 15731, blonde: 39397, light brown: 141414, dark brown: 127980, black: 14526, other: 4186, do not know/prefer not to answer 650. Individuals with missing data (“Prefer not to answer”, “Other”, “Do not know”) were excluded from all the analysis. For the red hair colour versus brown and black hair colour, the blonde individuals were removed (red=15731, non-red=283920), and self-reported red hair individuals were removed from the blonde versus brown and black analysis (blonde=39397, non-blonde=283920). For the comparison of brown versus black, light and dark brown individuals were combined and compared to black hair individuals (brown=269394, black=14526).

### 4. Genome-wide association and conditional analysis

Following individual and genotype-level QC a logistic regression model was used to regress presence/absence of each hair colour on bi-allelelic variant genotype assuming a (log) additive model, adjusting for the first 15 axes of variation from the PCA, and genotyping batch (equation [1]).

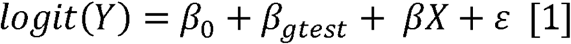

Conditional analyses used the same multivariate logistic regression model above, with the addition of the lead SNP (denoted by *βA*_glead_) from each signal of association [2].

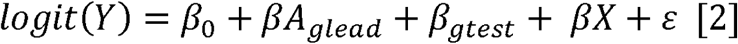

For chromosomes where more than one signal of association was apparent (lead SNP p ≤ 5×10^−8^), subsequent rounds of conditional analysis were performed, adding each new lead SNP as an extra conditioning SNP (denoted by *βA_glead_*, where *A_glead_* in equation [2] is a matrix of individual by lead SNP genotypes). To accelerate this stepwise procedure, we first removed the SNPs with p-value>0.1 in the initial GWAS, and then performed the conditional analysis until no single SNP association exceeded p ≤ 5×10^−8^. Finally, to check none were missed, SNPs with an initial p-value > 0.1 were included again and the conditional analysis continued until no SNPs exceeded p ≤ 5×10^−8^. There were just 3 SNPs that were ~10^−8^ in blonde hair colour, and only one extra round of conditional analysis was needed. Regression analyses were performed using PLINK v1.9. Manhattan and Q-Q plots were generated using the R package *qqman*^2^ and *ggplot2*.

### 5. Hair colour genetic scoring

To create a polygenic risk score for blonde hair, all lead variants associated with blonde hair were included into a least absolute shrinkage and selection operator (LASSO) model.

Regularized effect sizes were estimated by penalized logistic regression using the *glmnet* package in R^3^ and the degree of regularization was defined by the most regularized model within 1 standard error of the minimum. The regularized effect estimates were used to create genetic scores for blonde hair colour, taken as a linear combination of the regularized effect estimate weighted by the number of alleles carried (assuming an additive genetic model in all cases).

Non-British (n=44595) and related (n=64571) individuals were taken as independent test for the blonde hair genetic scoring. The summed genetic score for each individual was calculated using Plink v1.9.

### 6. Penetrance

The penetrance of allelic combinations was calculated by summing the number of individuals of a particular hair colour with a given allelic combination over the sum of all individuals (all hair colours) with that allele combination. The calculation and plot were made using the .ped and .map plink files, and ggplot2.

### 7. Epistasis

Variants within 3.0Mb of the index SNP (±1.5Mb either direction) for each signal of association were selected for epistasis analysis. Explicit testing of interactions between *MC1R* non-synonymous variants and hair colour associated loci was performed by comparing the following full [3] and reduced [4] logistic regression models:

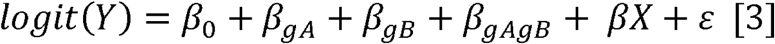

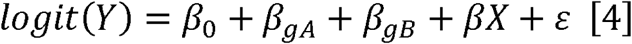

Where β_0_ is the model intercept, *β_gA_* is the regression coefficient for SNP A, *β_gB_* is the regression coefficient for SNP B and *β_gAgB_* is the joint regression coefficient for SNP A and B, i.e. the interaction term. Covariates were included in the model, denoted by the model term *βX*. For instance, population structure confounding may be accounted for by including N principal components in the model as covariates. Significance testing was performed using a likelihood ratio test, comparing model [3], with the reduced model [4] that omitted the interaction term, *β_gAgB_*. Epistasis testing was performed using *cassi* (see URLs) using the *--lr* command. SNP alleles that co-occur on the same haplotypes, but are in imperfect linkage disequilibrium, may generate the false impression of interactions, Wood et al.^4^ suggested including the main effects as covariates to remove these false interactions. Therefore, in the SNPs where there was apparent epistasis in chromosome 16, we corrected for rs34357723 and the MC1R SNPs, and if there were still signs of epistasis, we corrected for all the significant SNPs in the red versus brown and black hair conditional analysis. Correcting for these SNPs we removed most of the epistasis signals in chromosome 16.

### 8. Probabilistic inference of causal SNPs

We implemented the probabilistic inference of causal SNPs (PICS) approach^5^ to identify the best candidate SNP at a locus, taking into account the strength of association of the lead SNP and the LD with each SNP in the fine-mapping interval. PICS calculates the conditional posterior probability *P*(*B*^causal^ | *A* ^lead^), that is the probability that SNP B is the causal variant given SNP A is the lead SNP. Under the assumption that effect sizes for non-causal SNPs are drawn from a Normal distribution, *N*(*μ,σ*^2^), Farh *et al* derive from empirical data the sample standard deviation, *σ_s_*, and the expected mean association signal, *μ_s_*, which scales linearly with the LD to the true causal SNP.

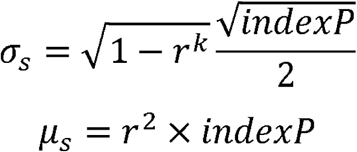

Where *indexp* is the *–log*_10_*P* of association of the causal SNP for that locus, that is taken to be the lead SNP for that fine-mapping interval. The calculated posterior probabilities were used to construct 95% credible intervals, defining the resolution of fine-mapping over a given genomic interval.

### 9. eQTL analysis

To test for the potential of non-coding association signals to impact on gene regulation we utilised expression quantitative trait loci (eQTL) summary statistics from the GTex^6,7^ and MuTHER^8,9^ studies in tissues relevant to pigmentation (MuTHER; skin biopsies, GTex; sun-exposed and sun not-exposed skin tissue). Summary statistics for genes that fell within 3Mb intervals of the lead SNP of each association signal were extracted. We wished to test the explicit hypothesis that the same causal variants underlie each eQTL and our association signal for each hair colour (H_A_). Subsequently the null hypothesis is a compound null of no association for either trait (H_01_), and the combinations of overlapping but independent eQTL and trait associations (H_02_) and an association in one trait/eQTL but not the other (H_03_ & H_04_). A Bayesian testing framework that tests the set of hypotheses relating to those defined above (H_01-4_ & H_A_) was developed by Giambartolomei *et al*, and is implemented in the R package *coloc*^10^. Subsequently we report the posterior probability for each association signal-gene eQTL signal and hypotheses H_02-04_ (PP1, PP2 & PP3) and the alternative hypothesis H_A_ (PP4).

Genotype and normalised expression data from GTEx were used to plot the ASIP eQTL (rs6059655).

### 10. Chromatin enrichment

#### Roadmap Epigenome, FANTOM5 and ENCODE

BED format files were downloaded for epigenetic marks associated with activated or accessible chromatin from the Roadmap Epigenome and ENCODE project websites (October 2017). Files were downloaded for annotations in primary melanocytes, keratinocytes, and fibroblasts included in the Roadmap Epigenome Project: H3K4me1, H3K4me3, H3K27ac and DNase I hypersensitive sites (DHS; where available). Primary melanocyte, keratinocyte and other pigmentation cell type transcriptional start site (TSS±100bp) data were downloaded from the FANTOM5 website (October 2017).

#### MITF ChIP-seq peak calling

Micropthalmia-associated transcription factor is generally considered the master regulator of melanogenesis in melanocytes. Short read data were downloaded from SRA corresponding to MITF ChIP-seq data derived from primary melanocytes and the melanocytic cell line Hermes3A^11–13^. Short sequencing reads were extracted using *fastq-dump* and assessed for quality using *fastQC* (see URLs). Reads were processed prior to genome alignment to remove adaptor contamination using *trimmomatic*. Processed reads were aligned to GRCh37/hg19 using BWA-aln with the options *-l 21 -k 2 -n 0.05 -o1* as described above. Peaks were called using MACS2 with an FDR 1% compared to a single common input^14^.

#### Annotation enrichment testing

For each cell type and annotation where more than one sample existed, intervals were merged using *Bedtools intersect* for two files and *multiIntersectBed* for > 2 files per cell type^15^. To test for genomic enrichment we used a method that tests for the local enrichment of lead SNPs and those in LD with genomic features by randomly shifting annotations to generate a null distribution; implemented in *GoShifter*^16^. LD values calculated from 1000Genomes phase 3 EUR reference panel were used in conjunction with the lead variant from each independent signal of association (after conditional analyses). Genomic annotations were provided as BED format for all annotations in all cell types described above. *GoShifter* was run for 10000 permutations, and enrichment p-values were adjusted for multiple testing using the Benjamini & Hochberg procedure implemented in the R function *p.adjust* with a 5% FDR.

## Supplementary Figure Legends

**Supp Figure 1.** QQ plots for the GWAS. Plots for red versus black plus brown hair colour, blonde versus black plus brown hair colour and black versus brown hair colour.

**Supp Figure 2.** Epistasis between MC1R variants and variants elsewhere, MC1R variants are as indicated, and loci elsewhere in the genome.

**Supp Figure 3.** Polygenic Phenotype Scores. Genetic scores derived from all lead variants from blonde versus brown plus black hair colour, assuming an additive genetic model, applied to confirmation cohorts within Biobank: 3^rd^ degree relatives or closer individuals and white non-British indivduals.

## Supplementary Tables

**Supplementary Table 1:** Matrix of penetrance of red hair phenotype for each genotype combination

**Supplementary Table 2:** Red hair vs black plus brown GWAS, including analysis following serial conditioning for successive chromosome 16 variants

**Supplementary Table 3:** Blonde hair vs black plus brown GWAS, including analysis following serial conditioning for variants chromosome by chromosome

**Supplementary Table 4:** Brown hair vs black GWAS, including analysis following serial conditioning for variants chromosome by chromosome

**Supplementary Tables 5-10:** eQTL data from GTex of skin gene expression; locus is the location of the variant, gene_name is the affected gene. Supplementary Table 5, red hair associated variants in sun-exposed skin. Supplementary Table 6, red hair associated variants in non-sun-exposed skin. Supplementary Table 7, blonde hair associated variants in sun exposed skin. Supplementary Table 8, blonde hair associated variants in non-sun-exposed skin. Supplementary Table 9, brown hair associated variants in sun exposed skin. Supplementary Table 10, brown hair associated variants in non-sun-exposed skin.

**Supplementary Tables 11-13:** eQTL data from TwinsUK Multiple Tissue Human Expression Resource of skin gene expression. Locus is the location of the variant, gene_name is the affected gene. Supplementary Table 11 red hair associated variants. Supplementary Table 12 blonde hair associated variants. Supplementary Table 13 brown hair associated variants.

**Supplementary Table 14.** Human candidate genes from blonde GWAS and mouse orthologues

**Supplementary Table 15.** Mouse orthologues of blonde candidate genes expressed in skin or skin appendages

